# Age-related development in prefrontal-subcortical resting-state functional connectivity in nonhuman primates

**DOI:** 10.1101/2023.07.19.549741

**Authors:** Harshawardhan U. Deshpande, Stephen J. Kohut

**Affiliations:** Behavioral Neuroimaging Laboratory, McLean Hospital and Department of Psychiatry, Harvard Medical School, Belmont, MA, USA

**Keywords:** nonhuman primate, functional brain connectivity, awake, adolescent, development, fMRI, cortical-subcortical

## Abstract

**Introduction:** Understanding age-related changes in functional connectivity (FC) with regards to the maturation trajectories of cortical-subcortical circuits is critical for identifying biomarkers for disease vulnerability. The present study investigated resting-state FC in adolescent and adult nonhuman primates to characterize developmentally-sensitive functional brain circuits.

**Methods:** Resting-state fMRI data were acquired in adolescent (33.3±1.59 months; n=22) and adult (117.29±2.86 months; n=15) squirrel monkeys and FC was compared in seven prefrontal and ten subcortical regions-of-interest (ROIs). The effect of subject age on FC between each pair of ROIs was evaluated to identify nodes with the most age-sensitive connections (hubs) which were then used in seed-to-whole-brain FC analyses. A subset of adolescents (n=7) was also assessed over 3 longitudinal scans to track changes in hub connectivity throughout adolescence.

**Results:** A significant effect of age on ROI-ROI FC was found for adolescent (p<0.001), but not adult, subjects (p=0.8). Evaluation of parameter estimates (β) for each ROI-ROI pair found three within-prefrontal (dorsolateral (dlPFC), dorsomedial (dmPFC), and medial orbitofrontal cortices), two within-subcortical (R amygdala and L hippocampus), and three between prefrontal-subcortical (dlPFC, dmPFC, L caudate) hubs with the highest number of age-related connections. Large-scale organizational differences were also observed between the adolescent and adult groups. Longitudinal scans found within-subject changes in FC consistent with group effect.

**Conclusions:** The relationship between changes in FC and age during adolescence indicates dynamic maturation of several prefrontal–subcortical circuits in nonhuman primates. These findings provide specificity in our understanding of the development of functional brain circuits during and into late adolescence.

**Significance:** Adolescence marks a period of rapid development in the brain, but also increased vulnerability to mental health disorders. Age-related prefrontal-subcortical resting-state functional connectivity was evaluated in awake adolescent and adult squirrel monkeys. Identification of functional connectivity differences highlight a network of hubs with a high number of connections evolving from early to late adolescence, indicating selectivity in maturation during different stages of aging. Compared to adults, adolescents also show several large-scale organizational differences in circuits originating from important seed regions-of-interest. Longitudinal analysis reveals functional connectivity trajectories emerging from early adolescence and maturing into adult-like patterns during late adolescence. These findings identify functional connections that change dramatically during adolescence suggesting specific circuits that could be at heightened sensitivity to disease vulnerability.

## 1. Introduction

Adolescence is a critical period marked by granular changes in behavioral and neural development that persist into adulthood. Aside from the prenatal period, the most substantial modifications in neural architecture occur during adolescence with several rapid changes in the structure and function of the brain leading to large-scale physical and mental maturation from early to late adolescence (1). While these changes may govern increased capacity for higher-order social, emotional, and cognitive processing that set the stage for healthy behavior in adulthood (2, 3), they also represent a vulnerable period in which environmental, pharmacological, and/or social influences may alter the trajectory of brain development, increasing risk for psychiatric/developmental disorders (4). Identifying age-related changes in functional pathways between brain regions within the dynamic period of adolescence is crucial to understand normative development in the underlying functional architecture and to develop novel interventions and treatment strategies for psychiatric disorders.

Non-invasive methods such as functional magnetic resonance imaging (fMRI) provide a means for systematically investigating the functional architecture of the brain through resting-state functional connectivity (rsFC). rsFC analyses capitalize on spontaneous fluctuations in the blood oxygen level dependent (BOLD) contrast in the brain at rest; regions in the brain that are anatomically and spatially distinct can be functionally linked through synchronous spontaneous BOLD fluctuations, providing highly reproducible networks of brain regions acting in implicit synchronicity in the absence of an explicit task (5). Previous studies have used fMRI to examine developmental changes in FC during human brain development. For instance, FC has been shown to increase between, and weaken within, resting-state networks (RSNs) as a function of age, a pattern that suggests consolidation of FC centered on large scale networks or network hubs such as dorsal attention, saliency/ventral attention, and somatomotor networks (6). Further, FC between frontal and subcortical regions appear to strengthen from childhood to adulthood (7, 8). A recent study combined rsFC data from several large-scale datasets (e.g., UK Biobank, Adolescent Brain Cognitive Development, Human Connectome Project, and the Developmental Chronnecto-Genomics) to characterize brain state adaptations in human adolescent and adult subjects, showing greater strengthening, modularity, and integration of the brain’s functional connections beyond adolescence (9). Consequently, rsFC has been proposed as a tool to identify biomarkers associated with risk for psychiatric disorders such as autism spectrum disorder (10), adolescent-onset schizophrenia (11), and depression (12) among others.

The majority of studies investigating rsFC development have grouped individuals into broad categories that encompass relatively wide age ranges – i.e., children, early or late adolescence, adults etc. Considering the heterogeneity in brain developmental trajectories of brain volume (13), cortical thickness (14), functional organization (15, 16), and behavioral patterns, examining whole-brain FC changes across adolescence may provide important information about the temporal dynamics of FC between key brain regions. While studies in human populations such as these have undoubtedly increased our understanding of the adolescent brain, limitations such as the diversity arising from environmental and genetic backgrounds, and the lack of precise control of experimental design often limit causal inferences (17). Nonhuman primates (NHP) have become increasingly important for understanding normative and disrupted brain function and connectivity. In addition to their behavioral, genetic, phylogenetic, and neurobiological similarity to humans (18, 19), NHPs can be studied under carefully controlled conditions and behavioral histories that are difficult to obtain in humans. The lifespan of NHPs also provides an experimentally tractable timescale to conduct longitudinal neurodevelopmental analyses which has advantages to provide greater statistical power (75) over other species, such as rodents, which have significantly shorter adolescent periods (i.e., days or weeks). Furthermore, neuroimaging studies have demonstrated that the architecture of NHP brains is similar to humans (20–22) providing a translationally relevant opportunity to conduct well-controlled longitudinal analyses of brain rsFC during developmental periods.

The present study sought to address these gaps by comparing rsFC measurements between samples of 22 adolescent and 15 adult squirrel monkeys intensively trained for awake fMRI. Seed-to-whole brain FC measurements were obtained through regions-of-interest selected based on their associations with cognitive and behavioral development in prior studies. The adolescent and adult groups were independently modeled to interrogate the effect of increasing age on ROI-ROI FC patterns. Finally, a cohort of seven adolescent subjects was studied longitudinally for changes in seed-to-whole-brain FC patterns over a period of eight months.

## 2. Methods

### 2.1 Subjects

Thirty-seven squirrel monkeys (*Saimiri sciureus*) served as subjects and comprised two groups: adolescents (mean 33.42 ± 1.52 months; *n*=22; 12 male/10 female) and adults (mean 117.29 ± 2.86 months; *n*=15; 6 male/9 female). The experimental protocol was approved by the Institutional Animal Care and Use Committee at McLean Hospital in a facility licensed by the US Department of Agriculture in accordance with guidelines provided by the Committee on Care and Use of Laboratory Animals of the Institute of Laboratory Animals Resources, Commission on Life Sciences. The housing and feeding procedures were identical to those described previously (22). The adolescent group were also subjects in a larger longitudinal study; all experiments were conducted in drug- and experimentally naive subjects.

### 2.2 Acclimation for Imaging Procedures

Please refer to the Supplement for details and (22).

### 2.3 MRI Acquisition parameters and processing

Details of the standard preprocessing pipeline are described in the Supplement. Because head motion can lead to an overestimation of short-distance FC and an underestimation of long-distance FC calculations (3, 23). Several steps (described in Supplement) were utilized to limit the influence of motion including extensive experimental acclimation, exclusion criteria, regression of motion parameters and derivatives, and inclusion of mean framewise displacement (FD) as a nuisance covariate in analyses.

### 2.5 Selection of ROIs

Seventeen prefrontal and subcortical regions-of-interest (ROIs) were selected based on their involvement in prior developmental studies in rodent, nonhuman primate, and human studies as well as their role in a range of cognitive and behavioral processes (24–27). Seven subdivisions were based on a functional parcellation of the squirrel monkey prefrontal cortex (ventromedial prefrontal cortex, vmPFC; dorsolateral prefrontal cortex, dlPFC; bilateral ventrolateral prefrontal cortex, R and L vlPFC; dorsomedial prefrontal cortex, dmPFC; medial orbitofrontal cortex, mOFC; and lateral orbitofrontal cortex, latOFC) (28). Ten subcortical ROIs (bilateral amygdala, putamen, caudate, hippocampus, and ventral striatum) were based on previously described anatomical landmarks (29–31).

### 2.6 ROI-ROI FC analyses to examine the effect of age in adolescent and adult monkeys

A mask dataset combining and labeling the seventeen ROIs was calculated (*AFNI: 3dCalc, 3drefit*). The effect of subject age (in months) on connectivity between the timeseries of the selected seventeen ROIs was assessed using a multivariate general linear modeling (MVM) framework. Subject-specific time series for each ROI were calculated (*AFNI: 3dNetCorr*) and analyzed for Fisher Z-transformed Pearson’s correlation connectivity estimates between each ROI pair using a wrapper to AFNI’s 3dMVM tool, provided as a part of the FATCAT toolbox (32). Subject age (months), sex, and motion (averaged FD) were included as covariates in the model along with the Fisher Z-transformed Pearson’s correlation connectivity datasets for each subject. Independent models were constructed for the adolescent and adult groups.

The multivariate model described here allows the treatment of correlations between each pairwise combination of ROIs as within-subject repeated-measures factors. Among MRI analysis software packages, the FATCAT MVM model uniquely adjusts for the effect of continuous covariates (33). If the main effect in the omnibus model of connectivity for any covariate is significant, it can justify the subsequent examination of the results of the *post hoc* models for each pairwise comparison. Specifically, the *post hoc* models determine whether the covariate is significant for each pairwise comparison. The parameter estimate (β) from these *post hoc* tests would also determine the nature of the correlation between age and FC, with a positive β indicating an increase in FC with an increase in age and a negative showing an inverse relationship. To visualize the parameter estimates (β) and p-values from ROI-ROI pairs, Z-scored FC from the ROI-ROI pairs were extracted for individual subjects (*AFNI: 3dROIstats*) and correlated with subject age (months) (*R: ggplot, cor*).

### 2.7 Classification of Hubs

The ROIs which were part of pairwise connections identified as contributing the most to the main age effect in the model described in section 2.6 were ranked based on the number of significant ROI pairs to which they belong. The ROIs with the highest number of connections were classified as within-PFC, within-subcortical, and between PFC-subcortical hubs.

### 2.8 Seed-to-whole-brain FC analyses

Using each of the identified hub ROIs as seeds in independent analyses, seed-to-whole-brain Z-scored FC maps were calculated for each individual (*AFNI: 3dNetCorr*). For each seed, group maps were compared for differences (adolescent vs. adults, *AFNI: 3dMVM*) with sex and mean FD included as covariates in the model. The minimum voxel cluster size for all whole-brain analyses was determined (*AFNI: 3dFWHMx, 3dClustSim*) using a two-component measure of the spatial autocorrelation of the preprocessed data (34). Maps were thresholded at a voxel-wise p=0.001, FDR-corrected p=0.05 (False Discovery Rate;(35)), and a cluster threshold of 20 voxels.

### 2.9 Longitudinal analyses of adolescent subjects

A subset of subjects (n=7) underwent fMRI scans with resting-state data collected thrice over a period of eight months. This particular subset of adolescents remained drug naive while the others (n=15) were subjects in adolescent drug exposure studies. The first scan was considered as the baseline and the second and third scans occurred six and eight months after the baseline scan respectively, providing a longitudinal analysis of normative brain FC development during adolescence. A multivariate model (*AFNI: 3dMVM*) was constructed for each of the hub ROIs identified in 2.7 with scan session, sex, and motion as covariates to examine the effect of scan session on seed-to-whole-brain FC, i.e., changes in FC from baseline to six months and then eight months due to normative brain aging. Maps were thresholded at a voxel-wise p=0.001, FDR-corrected p=0.05, and a cluster threshold of 20 voxels. For any clusters passing the threshold, subject-level FC Z-scores were extracted (*AFNI: 3dROIstats*) to visualize the FC change over the three scans.

## 3. Results

### 3.1 Subject metrics

The adolescent NHP group included 22 subjects (age range: 30-36 months) while the adult group included 15 subjects (age range: 109-121 months). The average FD was 0.14 (±0.11) mm for adolescents and 0.19 (±0.10) mm for the adults with no significant differences between the groups (p=0.18).

### 3.2 ROI-to-ROI FC relationship with age

The nature of the relationship between increasing age on FC between ROI-ROI pairs was investigated using the 17 ROIs described in section 2.5. The two groups were independently assessed for a relationship between ROI-ROI FC and subject age using the multivariate model described in section 2.7. The omnibus F-tests for sex and motion both did not show a significant relationship with FC, however, a significant effect of age on ROI-ROI FC was found for adolescents (p<0.001), but not adults (p=0.93). *Post hoc* tests were examined to determine which ROI-ROI pairs contributed to the effect of age on FC in the adolescent group. The parameter estimates (β) for each ROI-ROI pair are shown for adolescents (lower triangular matrix) and adults (upper triangular matrix) in Figure 1A. Figure 1B shows two exemplar age vs. FC correlations: dmPFC to L amygdala (upper panel, post-hoc β = 0.1634, p = 0.0029 for adolescents, β = 0.0089, p = 0.7079 for adults) and L vStr to R vStr (lower panel, post-hoc β = -0.1248, p = 0.0002 for adolescents, β = 0.0036, p = 0.8309 for adults). Between the dmPFC to L amygdala pair, FC increases with age for the adolescents (+ve β) while for the adults, FC does not show a relationship with age. Between the L vStr to R vStr pair, FC decreases with increasing age for adolescents (-ve β), a pattern not seen in adults. A majority of significantly contributing connections with an inverse age-FC relationship were between subcortical regions. In both exemplars, FC for the adolescents above 33 months of age is not different from that in adults (p=0.42 for dmPFC to L amygdala pair and p=0.8 for the L vStr to R vStr pair).

**Figure 1.**
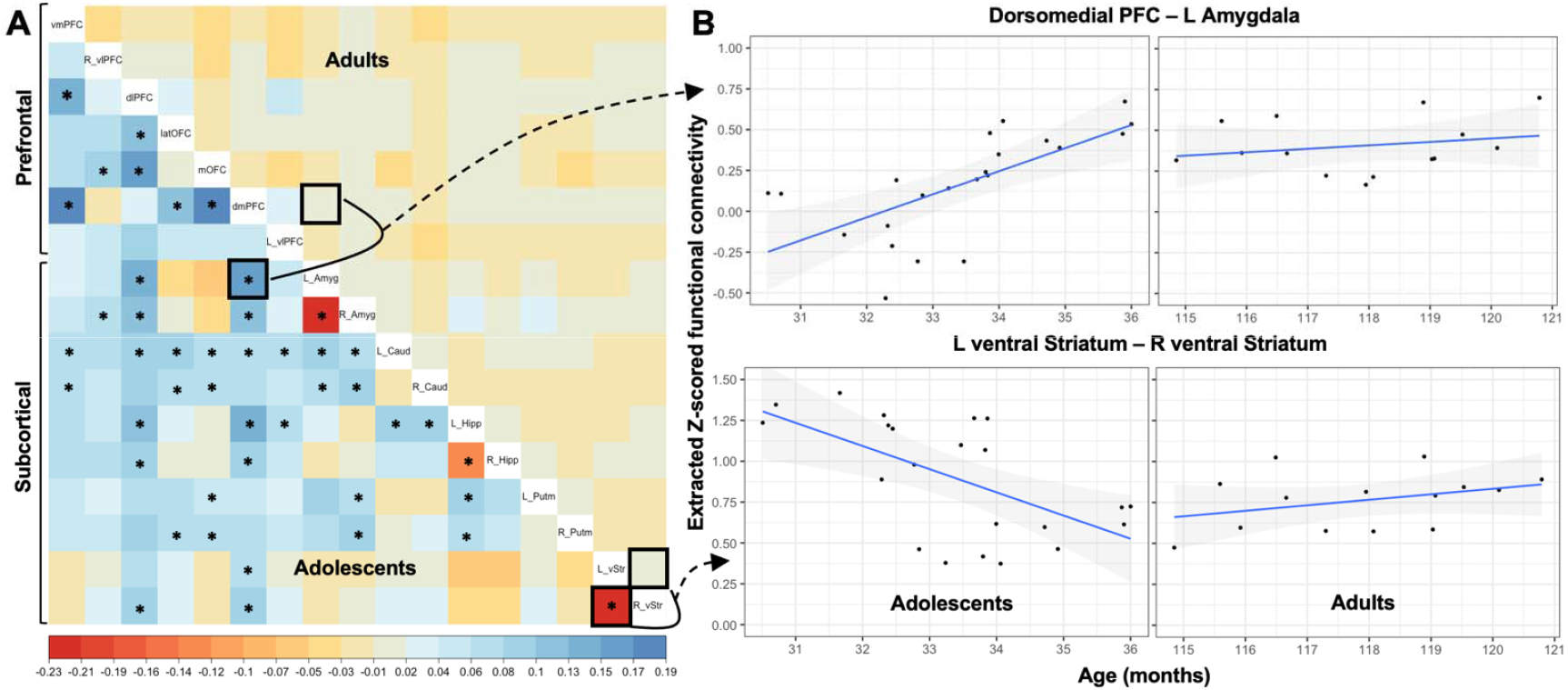
ROI-ROI functional connectivity models for the adolescent and adult groups. **A**. Parameter estimates (β) from the *post hoc* tests supporting the omnibus models of the effect of age on functional connectivity for the adolescent (n=22, lower triangular matrix) and adults (n=15, upper triangular matrix). Larger magnitudes of β-values indicates a stronger contribution of the ROI-ROI pair to the omnibus test and the polarity of β indicates the age-FC relationship directionality (+ve β = direct, -ve = inverse). Pairwise *post hoc* tests for which the adolescent group (lower triangular matrix) had p-values less than 0.05 are indicated with a *. **B**. Age-FC correlations for the dmPFC - L Amygdala (top, positive relationship) and L vStr - R vStr (bottom, inverse relationship) ROI pairs.

Figure 2 shows the ROI-ROI pairs with the strongest contribution to the omnibus test (high β, low p-values) for the adolescent group in the form of a circular plot. Among the prefrontal ROIs, the dlPFC, dmPFC, and the mOFC show a hub-like nature for an age-FC relationship with 3 pairwise connections to other prefrontal regions (Figure 2A). Within the subcortical ROIs, the R amygdala and L hippocampus are hubs with 5 pairwise connections to other subcortical regions (Figure 2B). Between the prefrontal and subcortical regions, the dmPFC, dlPFC, and L caudate are hubs with 5 connections each.

**Figure 2.**
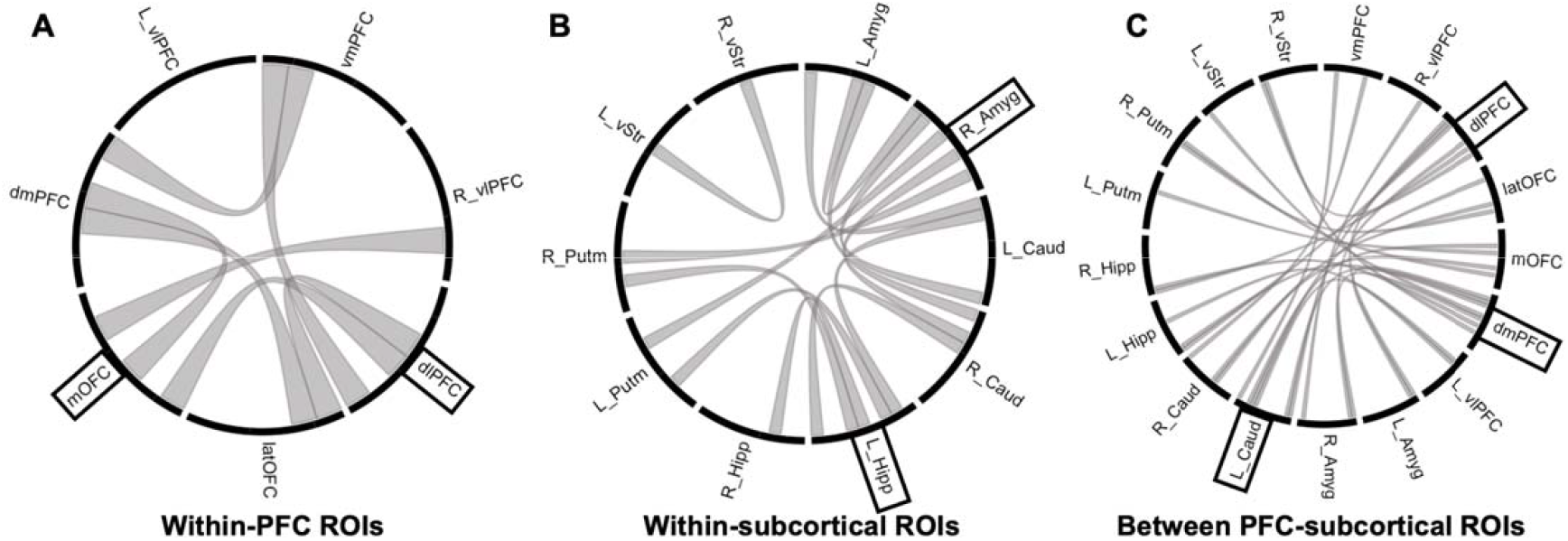
Circular plots showing most powerful (highest absolute β parameter estimates, lowest p-values) ROI-ROI pair connections contributing to the omnibus multivariate test within the PFC only (**A**), within subcortical ROIs only (**B**), and between the PFC and the subcortical ROIs (**C**). Among the seven PFC ROIs, the dlPFC and mOFC (3 connections) are the most connected. Within the ten subcortical ROIs, R Amygdala is a subcortical hub with 5 connections. The dmPFC (7), dlPFC (6), and the L Caudate (6 connections) can be identified as connectivity hubs between the PFC and subcortical ROIs.

### 3.3 Seed-to-whole-brain FC analyses: Adult group vs. adolescent group

#### 3.3.1 Prefrontal hubs as seeds

The three PFC regions identified as hubs (dlPFC, dmPFC, mOFC) were used as seeds to calculate seed-to-whole-brain FC maps comparing the adolescent and adult groups. The maps from each seed were thresholded (voxelwise-p = 0.001, FDR corrected p < 0.05, cluster threshold >= 20 voxels) to find regions with significant differences in FC between the groups. For the dlPFC and dmPFC (Fig. 3A and B), adolescents show greater connectivity with sensorimotor regions, attentional regions (posterior parietal, superior parietal cortices) while adults show greater connectivity with lateral septum and cerebellum. For the mOFC, adolescents show greater connectivity with posterior cingulate cortex while adults show greater connectivity with globus pallidus, septum, and dorsal hypothalamus. For the full list of regions, refer to Table 1.

**Figure 3.**
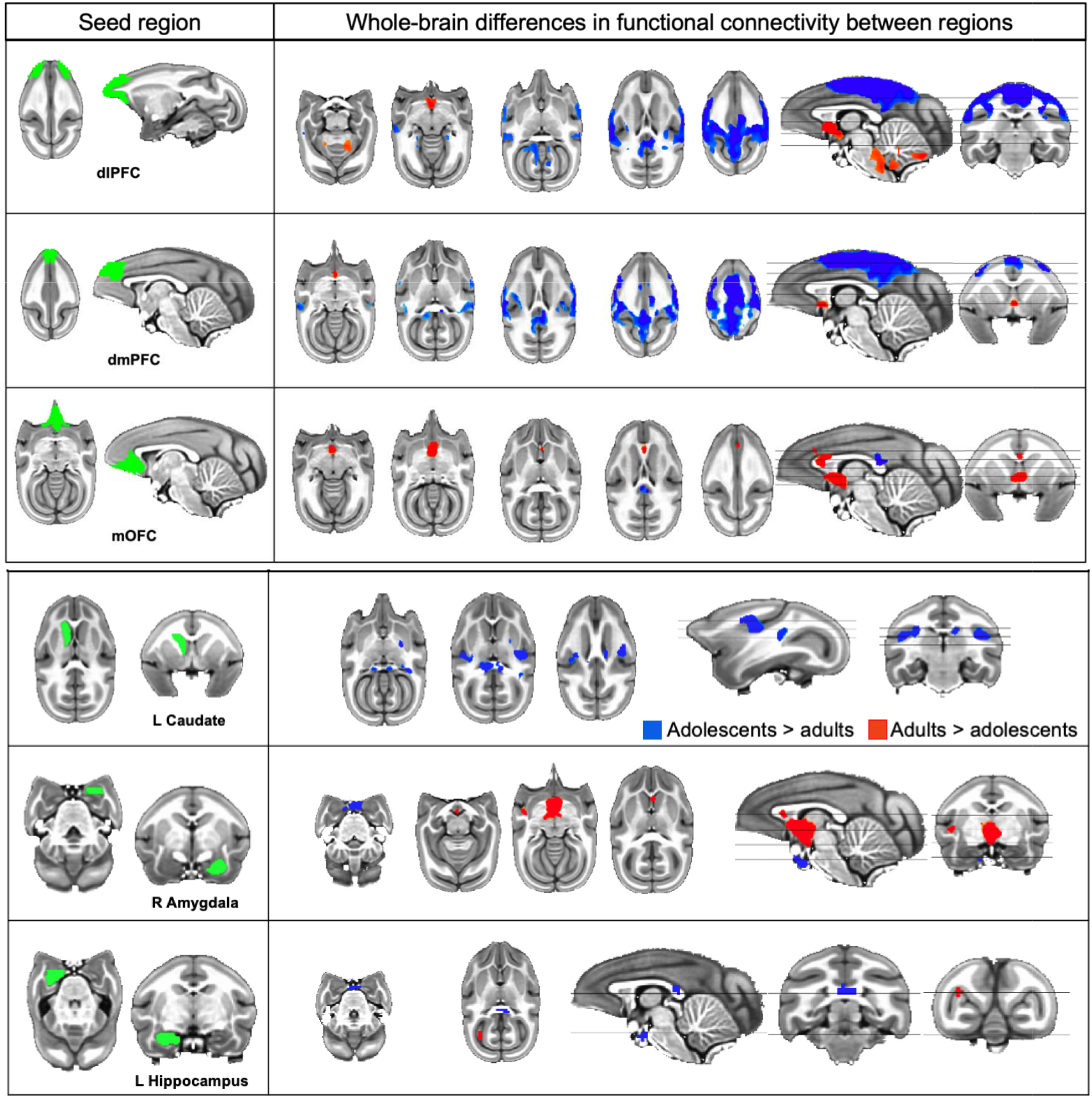
Seed-to-whole-brain functional connectivity differences between the adolescent and adult groups for hub regions. **A**. Blue (adolescents > adults) and red (adults > adolescents) regions with prefrontal hub ROIs (dlPFC, dmPFC, mOFC) as seeds (left column) in a seed-to-whole-brain FC analysis (p=0.001, FDR corrected p<0.05, minimum cluster threshold > 19 voxels). **B**. Blue (adolescents > adults) and red (adults > adolescents) regions with subcortical hub ROIs (L Caudate, R Amygdala, L Hippocampus) as seeds (left column) in a seed-to-whole-brain FC analysis (For L Caudate, R Amygdala maps, p=0.001, FDR corrected p<0.05, minimum cluster threshold > 19 voxels. For L Hippocampus map, no clusters survived FDR correction. Uncorrected clusters at p=0.001 shown). For connectivity maps for the other ROIs, see Supplement (Figure SF1 A-D, SF2 A-G).

**Table 1.**
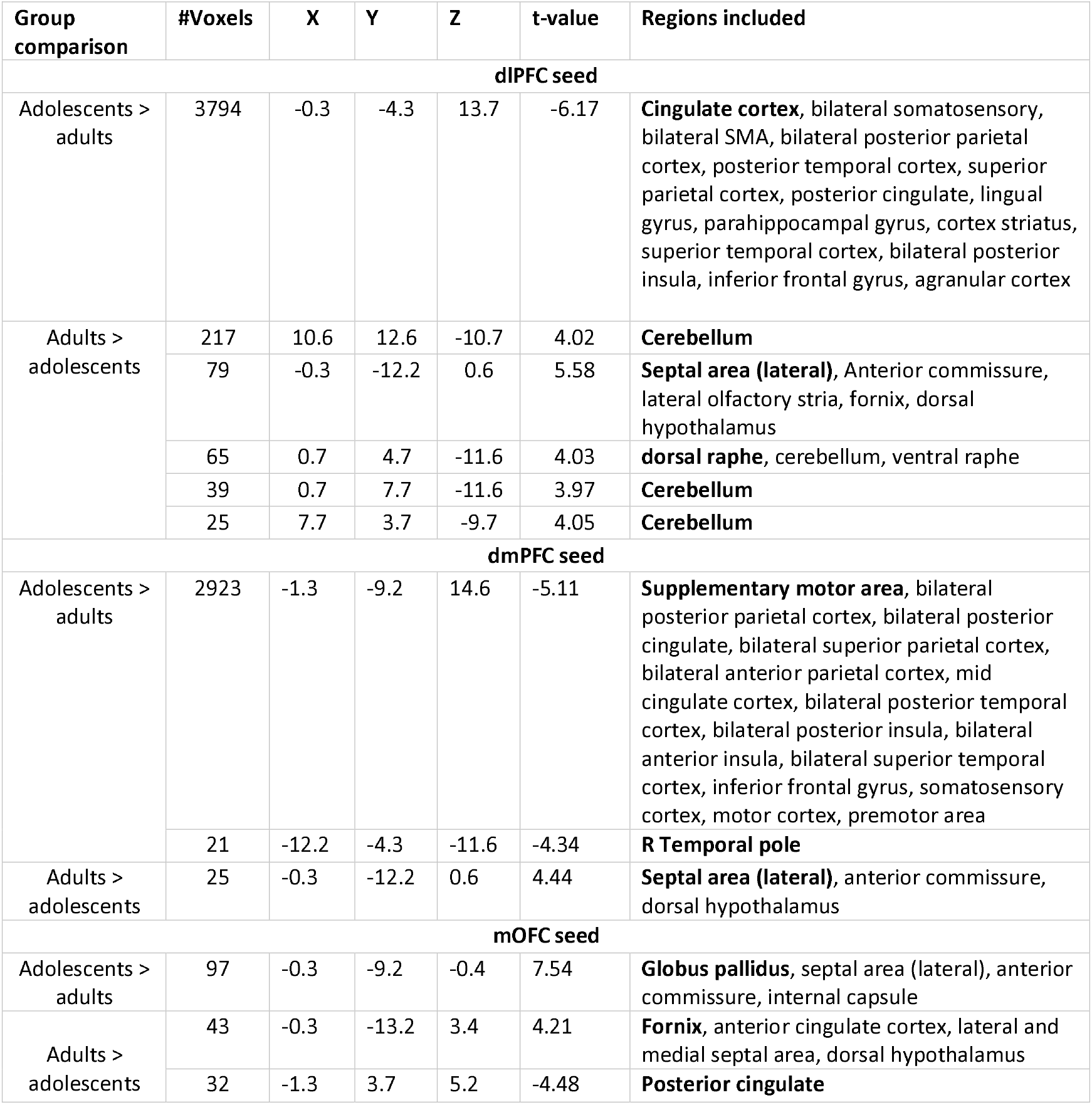
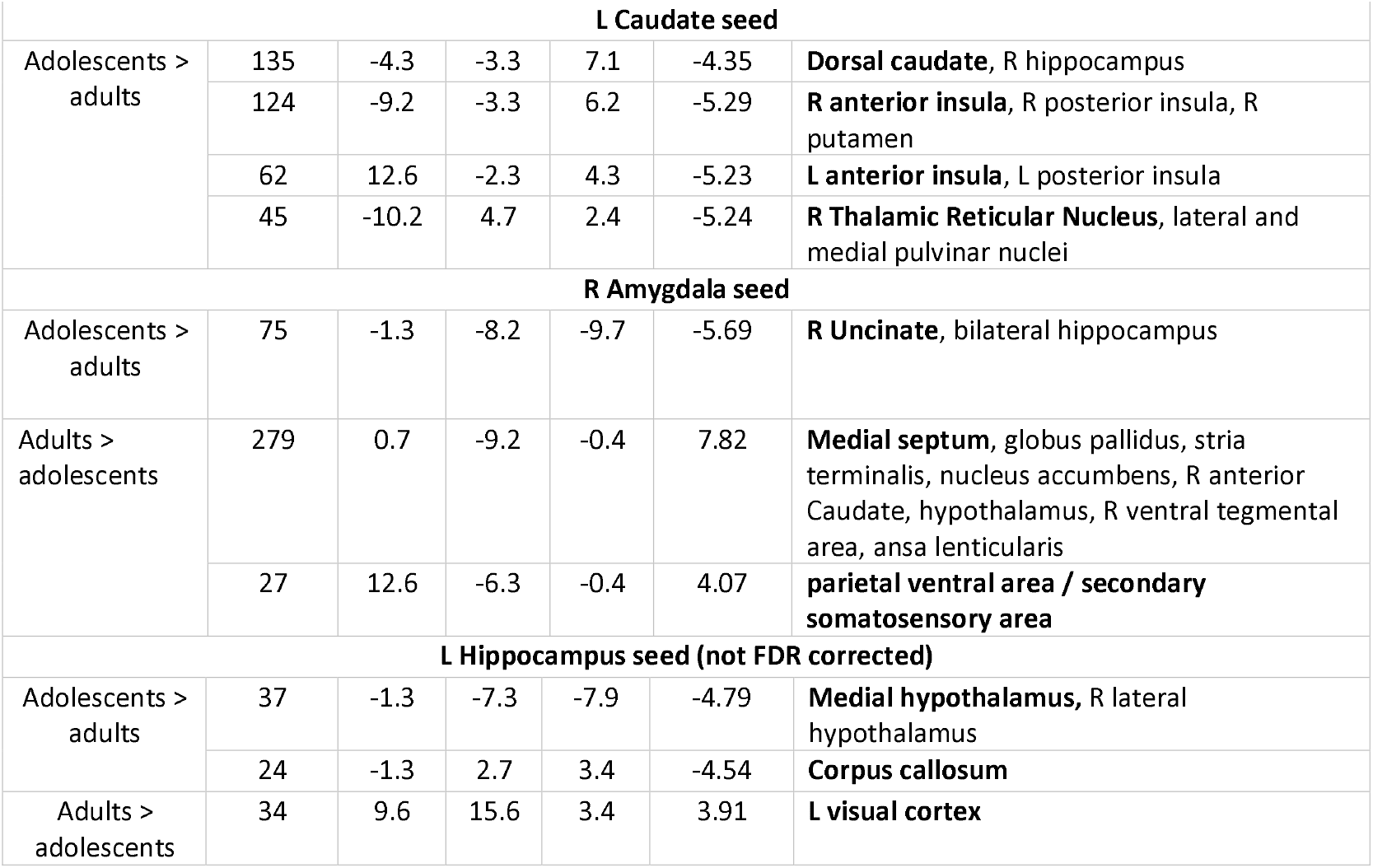
Seed-whole-brain functional connectivity differences for hubs identified in ROI analyses. For the six hubs identified, (dlPFC, dmPFC, mOFC, L Caudate, R Amygdala, L Hippocampus), the brain regions as identified using the squirrel monkey atlas are listed. Bolded region denotes the peak region and the X, Y, Z coordinates are listed for the peak region. For a list of regions for the other ROIs, see Supplement (Figure ST1, ST2).

| Group comparison | #Voxels | X | Y | Z | t-value | Regions included |
| --- | --- | --- | --- | --- | --- | --- |
| **dIPFC seed** |  |  |  |  |  |  |
| Adolescents > adults | 3794 | -0.3 | -4.3 | 13.7 | -6.17 | **Cingulate cortex**, bilateral somatosensory, bilateral SMA, bilateral posterior parietal cortex, posterior temporal cortex, superior parietal cortex, posterior cingulate, lingual gyrus, parahippocampal gyrus, cortex striatus, superior temporal cortex, bilateral posterior insula, inferior frontal gyrus, agranular cortex |
| Adults > adolescents | 217 | 10.6 | 12.6 | -10.7 | 4.02 | **Cerebellum** |
|  | 79 | -0.3 | -12.2 | 0.6 | 5.58 | **Septal area (lateral)**, Anterior commissure, lateral olfactory stria, fornix, dorsal hypothalamus |
|  | 65 | 0.7 | 4.7 | -11.6 | 4.03 | **dorsal raphe**, cerebellum, ventral raphe |
|  | 39 | 0.7 | 7.7 | -11.6 | 3.97 | **Cerebellum** |
|  | 25 | 7.7 | 3.7 | -9.7 | 4.05 | **Cerebellum** |
| **dmPFC seed** |  |  |  |  |  |  |
| Adolescents > adults | 2923 | -1.3 | -9.2 | 14.6 | -5.11 | **Supplementary motor area**, bilateral posterior parietal cortex, bilateral posterior cingulate, bilateral superior parietal cortex, bilateral anterior parietal cortex, mid cingulate cortex, bilateral posterior temporal cortex, bilateral posterior insula, bilateral anterior insula, bilateral superior temporal cortex, inferior frontal gyrus, somatosensory cortex, motor cortex, premotor area |
|  | 21 | -12.2 | -4.3 | -11.6 | -4.34 | **R Temporal pole** |
| Adults > adolescents | 25 | -0.3 | -12.2 | 0.6 | 4.44 | **Septal area (lateral)**, anterior commissure, dorsal hypothalamus |
| **mOFC seed** |  |  |  |  |  |  |
| Adolescents > adults | 97 | -0.3 | -9.2 | -0.4 | 7.54 | **Globus pallidus**, septal area (lateral), anterior commissure, internal capsule |
| Adults > adolescents | 43 | -0.3 | -13.2 | 3.4 | 4.21 | **Fornix**, anterior cingulate cortex, lateral and medial septal area, dorsal hypothalamus |
|  | 32 | -1.3 | 3.7 | 5.2 | -4.48 | **Posterior cingulate** |
| **L Caudate seed** |  |  |  |  |  |  |
| Adolescents > adults | 135 | -4.3 | -3.3 | 7.1 | -4.35 | **Dorsal caudate**, R hippocampus |
|  | 124 | -9.2 | -3.3 | 6.2 | -5.29 | **R anterior insula**, R posterior insula, R putamen |
|  | 62 | 12.6 | -2.3 | 4.3 | -5.23 | **L anterior insula**, L posterior insula |
|  | 45 | -10.2 | 4.7 | 2.4 | -5.24 | **R Thalamic Reticular Nucleus**, lateral and medial pulvinar nuclei |
| **R Amygdala seed** |  |  |  |  |  |  |
| Adolescents > adults | 75 | -1.3 | -8.2 | -9.7 | -5.69 | **R Uncinate**, bilateral hippocampus |
| Adults > adolescents | 279 | 0.7 | -9.2 | -0.4 | 7.82 | **Medial septum**, globus pallidus, stria terminalis, nucleus accumbens, R anterior Caudate, hypothalamus, R ventral tegmental area, ansa lenticularis |
|  | 27 | 12.6 | -6.3 | -0.4 | 4.07 | **parietal ventral area / secondary somatosensory area** |
| **L Hippocampus seed (not FDR corrected)** |  |  |  |  |  |  |
| Adolescents > adults | 37 | -1.3 | -7.3 | -7.9 | -4.79 | **Medial hypothalamus**, R lateral hypothalamus |
|  | 24 | -1.3 | 2.7 | 3.4 | -4.54 | **Corpus callosum** |
| Adults > adolescents | 34 | 9.6 | 15.6 | 3.4 | 3.91 | **L visual cortex** |

#### 3.3.2 Subcortical hubs as seeds

The three subcortical regions identified as hubs (R amygdala, L hippocampus, L caudate) were used as seeds to calculate seed-to-whole-brain FC maps comparing the adolescent and adult groups. The maps from the seeds were thresholded (voxelwise-p = 0.001, FDR corrected p < 0.05, cluster threshold >= 20 voxels) to find whole-brain regions with significant differences in FC between the groups. For R amygdala (Fig. 2D), adolescents show greater connectivity with bilateral hippocampus while the adults show greater connectivity with medial septum and hypothalamus. For L caudate (Fig. 2E), adolescents show greater connectivity with dorsal caudate, bilateral insula cortices and right hippocampus. For L hippocampus, adolescents showed greater connectivity with posterior cingulate tail while the adults showed greater connectivity with left visual cortex. For the full list of regions, refer to Table 1.

### 3.4 Longitudinal changes in hippocampus to whole-brain FC with increasing age

The seven subjects with three longitudinal scans were assessed for differences in seed-whole-brain FC for the 17 seeds between the three scan time points (baseline, +6-months, +8-months). For each of the hubs, the seed-to-whole-brain maps at the three sessions were compared. Difference between the +6-month scan and baseline (+6-months > baseline) was observed (e.g., main effect of session for dmPFC in Figure 4, voxelwise-p = 0.001, FDR corrected p<0.05, cluster threshold ≥ 20 voxels). The z-scored FC between the hub seeds and the brain regions with FC differences in session was extracted for all three scan time points. FC increases from baseline to the +6-month scan and then does not change significantly to the +8-month scan (Figure 4B).

**Fig. 4.**
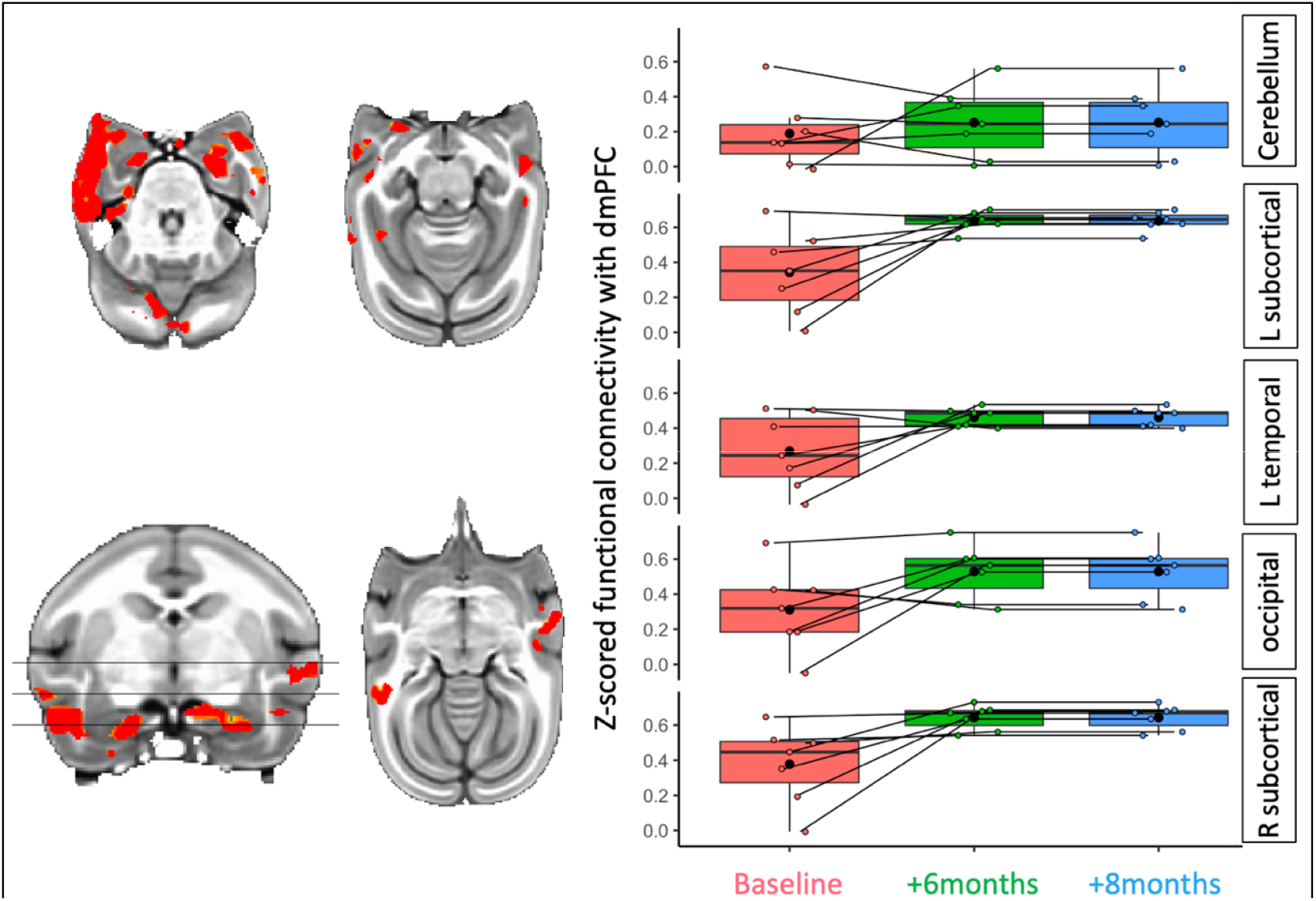
Longitudinal functional connectivity changes with the dmPFC. A. Main effect of the session (baseline, +6months, +8months) thresholded at p<0.001, FDR-corrected at p<0.05, cluster size > 40 voxels. B. ROI-extracts of z-scored functional connectivity for the three sessions between the dmPFC and the bilateral subcortical, L temporal, and occipital regions showing an increase from baseline to the +6 months session and then a leveling trend for the +8 months session. The control region (cerebellum) does not show such a trend.

## 4. Discussion

In the present study, we report age-related patterns of resting-state brain FC in a large sample of awake squirrel monkeys, a species with increasing relevance to the field of developmental neuroscience (17, 75). To do so, we leveraged a multivariate model to evaluate FC within and between brain regions previously associated with cognitive, social, and motor development and identified age-related changes in FC for several within-cortical, within-subcortical, and between PFC-subcortical connections in adolescent, but not adult subjects. These analyses also identified cortical and subcortical hub regions that showed the highest number of dynamic connections. Hubs were further evaluated using a seed-to-whole-brain approach to characterize brain-wide functional organizational differences between adolescents and adults. Finally, longitudinal FC changes in hub regions were tracked in a subset of adolescent subjects over an 8-month period. Collectively, these results show age-related reconfiguration of brain FC toward functional maturation specific to normative adolescents.

### PFC-subcortical FC changes with age within the adolescence period

The adolescent and adult groups were modeled independently to investigate the main effect of increasing age on changes in FC between 17 ROIs. All within-PFC and between PFC-subcortical connections increased as a function of age. In contrast, few (3 out of 13) within-subcortical connections showed a negative relationship, i.e., as age increased, FC decreased. These findings corroborate prior literature showing stronger PFC-PFC and PFC-subcortical connectivity with increasing age in humans. For example, Baum et al. (16) showed refinement in functional organization of the brain with greater modular segregation that corresponded with increased executive function as adolescents mature into adulthood. Vasa et. al (36) demonstrated increasing cortico-cortical and cortico-subcortical (e.g., between amygdala and frontal association cortices) FC over the course of adolescence. Recent work also found a negative relationship between brain regions with structural differentiation and FC, specifically, between sensorimotor regions, and default mode networks (37) highlighting structural brain network reconfiguration and the consequential functional maturation that occurs during adolescent development. Our findings suggest specific patterns of age-related brain FC reorganization that may be especially vulnerable to environmental perturbations during adolescent development and increased propensity for onset of mental health disorders.

### FC hubs showing age-related changes during adolescence

A typical feature of functional organization in the brain is that while most nodes have a limited number of connections, a small number are highly connected, forming so-called hubs (38). Previous studies investigating FC hubs in the human brain have noted that the strongest hubs are located in regions associated with the default mode network (i.e., dmPFC, hippocampus, posterior cingulate cortex, angular gyrus) and those associated with sensory function, while subcortical networks exhibit weaker hubs (39). Further, hub connections tend to favor local, rather than long range, connectivity in non-human primates (40), which has also been shown to be consistent with mouse (41). A study in awake marmosets (42) demonstrated prominent visual, thalamic and a DMN-like network hubs with widespread connectivity reflecting those found in the human brain. In our data, we identified several highly connected hubs at three levels of organization: 1) within-PFC: dmPFC, dlPFC, and mOFC; 2) within-subcortical: R amygdala and L hippocampus, and; 3) between PFC and subcortical regions: dmPFC, dlPFC, and L caudate.

### Within-PFC hubs

Several within-cortical networks are thought to work in concert to mediate cognitive functions contributing to behavioral control, emotional regulation, and decision-making (7, 43). Among the within-PFC local hubs identified here, mOFC is said to be a part of the orbital and medial prefrontal cortex (OMPFC)(44, 45) network and plays a role in decision making, monitoring the reward value of stimuli (46), and integrating somatosensory and multimodal stimuli (47) to guide behavior and decision-making based on expected outcomes (48). dlPFC works in coordination with OMPFC, the dorsal attention network, and the central executive network to support higher order cognitive processes such as inhibitory control, attentional control, planning, and problem-solving (49). Specifically, the dlPFC’s connectivity with: 1) lateral OFC facilitates the integration of cognitive and emotional information, and decision-making (50), 2) anterior cingulate cortex (ACC) to regulate attention by detecting errors and triaging competing cognitive processes (51), and 3) vmPFC for emotion regulation, social cognition, and self-referential processing (52). The overall pattern of FC strengthening between these PFC hubs and between the hubs and other PFC regions likely reflects the maturation of cognitive processes such as goal-directed behavior and decision-making.

### Within-subcortical hubs

Subcortical areas in the brain, while showing weaker hubs (39), comprise several RSNs (DMN, cingulo-opercular network) that support foundational capacities such as attention regulation and cognitive performance. As a local, within-subcortical hub, R amygdala is primarily involved with emotional processing and the regulation of emotional responses (53, 54). The second within-subcortical hub identified was the L hippocampus, which is often associated with memory formation, cognition, and emotional processing (55). The connections of these hubs with other subcortical structures such as L amygdala (integration of emotional information across hemispheres), R hippocampus (memory consolidation for emotionally salient events), and caudate (integration of emotional valence with reward-related processes) implicate these hubs as parts of networks crucial for the development of emotional and memory salience.

### Between PFC-subcortical hubs

Considering the between PFC and subcortical hubs, dlPFC and the dmPFC both showed greater reward-related connectivity associated with years since adolescent onset substance use (56). The intrinsic (resting-state) connectivity between dlPFC and subcortical regions (specifically thalamus) has been associated with the age-related changes in learning performance in adolescents and young adults (57). Additionally, dlPFC has also been shown to be connected with other subcortical regions such as hippocampus (integration of memory with executive functions), amygdala (integration of emotional processing with cognitive control and emotional regulation), and ventral striatum (reward processing, motivation, and goal-directed behaviors), supporting a range of cognitive, emotional, and motoric functions (58, 59). The attentional RSN comprising dlPFC has shown to be strongly connected to dorsal striatum (caudate and putamen) in both NHPs and humans (21). L caudate has been shown to be a major pathway between PFC and subcortical regions, through connections to the prefrontal and orbitofrontal cortex (decision-making, craving, reward processing), ACC (conflict monitoring), thalamus (attention regulation), hippocampus (spatial memory), and amygdala (emotional processing and fear response) in NHP (21) and human (60–62) studies. The PFC-subcortical hubs identified here form the cornerstones of a PFC-subcortical circuit, serving as integrators of information related to executive function, emotional regulation, and reward processing, and cognitive processes shown to develop during adolescence.

### Large-scale FC organizational differences between adolescents and adults

An investigation of differences between adolescents and adults using seed-to-whole-brain connectivity found that for the prefrontal hubs (dlPFC, dmPFC and mOFC), adolescents showed overall greater FC with somatomotor cortex, parietal attentional regions, posterior cingulate cortex while adults showed greater FC with the cerebellum and subcortical structures such as septum, fornix, hypothalamus. Recently, Rao et al. (2021) described diverse developmental patterns in rhesus macaque RSNs with fine-tuned changes from adolescence into early adulthood (63). Specifically, using a group IC analysis, they identified eight rhesus macaque RSNs and showed maturation of the sensorimotor network during childhood and protracted developmental trajectory of frontal network from childhood to early adulthood. Yassin et al. (2023) also found greater connectivity in PFC and cingulate networks to occipital and temporal regions, respectively, in adult compared to adolescent squirrel monkeys (22).

### Longitudinal changes in FC from early to late adolescence

These developmental patterns were also evident in the longitudinal analysis, indicating RSNs in squirrel monkeys that are conserved during adolescence which mature into adulthood. Compared to the baseline scan, adolescents FC increased during the +6 and +8-month scans, with connectivity strength approaching that in adults. Developmental and experiential synaptic pruning throughout adolescence has been shown to result in more efficient large-scale functional reorganization (integration and segregation) in human adolescence (64). Taken together, our findings corroborate prior work in humans showing the protracted development and refinement of complex cognitive processes that require simultaneous engagement of specialized brain regions facilitating proactive response planning and flexibility throughout adolescence (65)(66) and showing individual differences in connectivity predicting cognitive performance (67).

The methodological strengths of the current study include a large sample size covering a wide range of ages (adolescent and adult) encompassing the early to late adolescence period and a rapid longitudinal sampling of data facilitating the assessment of age-related changes. Our data were well-controlled for motion and were true resting fMRI scans (i.e., non-anesthetized). Recent work found few differences between adolescent and adult NHP using an ICA-based comparison of resting-state data (22); our fine-grained FC analyses identified several specific neural circuits associated with adolescent development, while also illuminating large-scale organizational differences between adolescents and adults. Our longitudinal approach allowed for the identification of true aging effects by capturing the development process as a within-subject effect and potentially allowing the investigation of individual variability over three time points covering the adolescence period (68). Together, these findings provide specificity in our understanding of the development of functional brain circuits during and into late adolescence. Future adolescent brain neuroimaging studies should also include regions discovered through whole-brain analyses such as insula, thalamus, ACC, PCC, and cerebellum to assess their developmental trajectories.

## Supporting information

Supplemental_material_Deshpande_Kohut

## Acknowledgements

We thank Jessi Stover, Julia Cunningham, Nora Monahan, Samantha McGouldrick, Bryan Carlson, and Craig Stone for their efforts in acclimating subjects to awake MRI procedures and Kenroy Cayetano for engineering the experimental equipment used for these studies. We also thank Drs. Paul Taylor, Gang Chen (National Institute of Mental Health – AFNI), and Tom Ross (National Institute on Drug Abuse) for their help in understanding and setting up some of the intricate multivariate modeling described in the paper. We are grateful to Dr. Rui Yuan, Carlo Servando De Los Angeles, and Dr. Vinod Menon for providing parcellation maps of the squirrel monkey prefrontal cortex. Finally, we thank Drs. Jack Bergman and Scott Lukas for intellectual discussion and feedback on the manuscript.

## REFERENCES

1. National Academies of Sciences, Engineering, and Medicine; Health and Medicine Division; Division of Behavioral and Social Sciences and Education; Board on Children, Youth, and Families; Committee on the Neurobiological and Socio-behavioral Science of Adolescent Development and Its Applications, The Promise of Adolescence: Realizing Opportunity for All Youth, E. P. Backes, R. J. Bonnie, Eds. (National Academies Press (US)).

2. B. J. Casey, A. Galván, L. H. Somerville, Beyond simple models of adolescence to an integrated circuit-based account: A commentary. Dev. Cogn. Neurosci. 17, 128–130 (2016).

3. J. D. Power, D. A. Fair, B. L. Schlaggar, S. E. Petersen, The development of human functional brain networks. Neuron 67, 735–748 (2010).

4. A. M. Gregory, et al., Juvenile mental health histories of adults with anxiety disorders. Am. J. Psychiatry 164, 301–308 (2007).

5. B. B. Biswal, et al., Toward discovery science of human brain function. Proc. Natl. Acad. Sci. U. S. A. 107, 4734–4739 (2010).

6. R. F. Betzel, et al., Changes in structural and functional connectivity among resting-state networks across the human lifespan. Neuroimage 102 Pt 2, 345–357 (2014).

7. S. Gu, et al., Emergence of system roles in normative neurodevelopment. Proc. Natl. Acad. Sci. U. S. A. 112, 13681–13686 (2015).

8. L. Q. Uddin, Salience processing and insular cortical function and dysfunction. Nat. Rev. Neurosci. 16, 55–61 (2015).

9. A. Abrol, et al., Developmental and aging resting functional magnetic resonance imaging brain state adaptations in adolescents and adults: A large N (>47K) study. Hum. Brain Mapp. 44, 2158–2175 (2023).

10. X. Yang, et al., A study of brain networks for autism spectrum disorder classification using resting-state functional connectivity. Machine Learning with Applications 8, 100290 (2022).

11. S. Wang, et al., Abnormal regional homogeneity as a potential imaging biomarker for adolescent-onset schizophrenia: A resting-state fMRI study and support vector machine analysis. Schizophr. Res. 192, 179– 184 (2018).

12. C. G. Connolly, et al., Resting-state functional connectivity of the amygdala and longitudinal changes in depression severity in adolescent depression. J. Affect. Disord. 207, 86–94 (2017).

13. K. L. Mills, et al., Structural brain development between childhood and adulthood: Convergence across four longitudinal samples. Neuroimage 141, 273–281 (2016).

14. N. Vijayakumar, et al., Brain development during adolescence: A mixed-longitudinal investigation of cortical thickness, surface area, and volume. Hum. Brain Mapp. 37, 2027–2038 (2016).

15. D. D. Jolles, M. A. van Buchem, E. A. Crone, S. A. R. B. Rombouts, A Comprehensive Study of Whole-Brain Functional Connectivity in Children and Young Adults. Cereb. Cortex 21, 385–391 (2010).

16. G. L. Baum, et al., Modular Segregation of Structural Brain Networks Supports the Development of Executive Function in Youth. Curr. Biol. 27, 1561–1572.e8 (2017).

17. J. Royo, S. J. Forkel, P. Pouget, M. Thiebaut de Schotten, The squirrel monkey model in clinical neuroscience. Neurosci. Biobehav. Rev. 128, 152–164 (2021).

18. R. M. Hutchison, S. Everling, Monkey in the middle: why non-human primates are needed to bridge the gap in resting-state investigations. Front. Neuroanat. 6, 29 (2012).

19. E. M. Weerts, B. J. Kaminski, Biological Research on Addiction: Chapter 4. Nonhuman Primate Models of Alcohol Abuse and Alcoholism (Elsevier Inc. Chapters, 2013).

20. D. S. Margulies, et al., Precuneus shares intrinsic functional architecture in humans and monkeys. Proc. Natl. Acad. Sci. U. S. A. 106, 20069–20074 (2009).

21. Y. Hori, et al., Altered Resting-State Functional Connectivity Between Awake and Isoflurane Anesthetized Marmosets. Cereb. Cortex 30, 5943–5959 (2020).

22. W. Yassin, et al., Resting state networks of awake adolescent and adult squirrel monkeys using ultra-high field (9.4T) functional magnetic resonance imaging. bioRxiv (2023) https://doi.org/10.1101/2023.01.08.523000.

23. K. R. A. Van Dijk, M. R. Sabuncu, R. L. Buckner, The influence of head motion on intrinsic functional connectivity MRI. Neuroimage 59, 431–438 (2012).

24. S. J. Lupien, B. S. McEwen, M. R. Gunnar, C. Heim, Effects of stress throughout the lifespan on the brain, behaviour and cognition. Nat. Rev. Neurosci. 10, 434–445 (2009).

25. C. A. Burghy, et al., Developmental pathways to amygdala-prefrontal function and internalizing symptoms in adolescence. Nat. Neurosci. 15, 1736–1741 (2012).

26. D. M. Demaster, S. Ghetti, Developmental differences in hippocampal and cortical contributions to episodic retrieval. Cortex 49, 1482–1493 (2013).

27. D. G. Gee, et al., Neurocognitive Development of Motivated Behavior: Dynamic Changes across Childhood and Adolescence. J. Neurosci. 38, 9433–9445 (2018).

28. R. Yuan, et al., Long-term effects of intermittent early life stress on primate prefrontal-subcortical functional connectivity. Neuropsychopharmacology 46, 1348–1356 (2021).

29. A. G. Lee, et al., Striatal dopamine D2/3 receptor regulation by stress inoculation in squirrel monkeys. Neurobiol Stress 3, 68–73 (2016).

30. D. M. Lyons, C. Yang, A. M. Sawyer-Glover, M. E. Moseley, A. F. Schatzberg, Early life stress and inherited variation in monkey hippocampal volumes. Arch. Gen. Psychiatry 58, 1145–1151 (2001).

31. D. M. Lyons, Stress, depression, and inherited variation in primate hippocampal and prefrontal brain development. Psychopharmacol. Bull. 36, 27–43 (2002).

32. P. A. Taylor, Z. S. Saad, FATCAT: (an efficient) Functional and Tractographic Connectivity Analysis Toolbox. Brain Connect. 3, 523–535 (2013).

33. G. Chen, N. E. Adleman, Z. S. Saad, E. Leibenluft, R. W. Cox, Applications of multivariate modeling to neuroimaging group analysis: a comprehensive alternative to univariate general linear model. Neuroimage 99, 571–588 (2014).

34. R. W. Cox, G. Chen, D. R. Glen, R. C. Reynolds, P. A. Taylor, FMRI Clustering in AFNI: False-Positive Rates Redux. Brain Connect. 7, 152–171 (2017).

35. C. R. Genovese, N. A. Lazar, T. Nichols, Thresholding of statistical maps in functional neuroimaging using the false discovery rate. Neuroimage 15, 870–878 (2002).

36. F. Váša, et al., Conservative and disruptive modes of adolescent change in human brain functional connectivity. Proc. Natl. Acad. Sci. U. S. A. 117, 3248–3253 (2020).

37. B.-Y. Park, et al., Adolescent development of multiscale structural wiring and functional interactions in the human connectome. Proc. Natl. Acad. Sci. U. S. A. 119, e2116673119 (2022).

38. M. P. van den Heuvel, O. Sporns, Network hubs in the human brain. Trends Cogn. Sci. 17, 683–696 (2013).

39. D. Tomasi, N. D. Volkow, Association between functional connectivity hubs and brain networks. Cereb. Cortex 21, 2003–2013 (2011).

40. R. Wu, et al., Graph theory analysis identified two hubs that connect sensorimotor and cognitive and cortical and subcortical nociceptive networks in the non-human primate. Neuroimage 257, 119244 (2022).

41. A. Liska, A. Galbusera, A. J. Schwarz, A. Gozzi, Functional connectivity hubs of the mouse brain. Neuroimage 115, 281–291 (2015).

42. A. M. Belcher, et al., Functional Connectivity Hubs and Networks in the Awake Marmoset Brain. Front. Integr. Neurosci. 10, 9 (2016).

43. D. R. Dajani, L. Q. Uddin, Demystifying cognitive flexibility: Implications for clinical and developmental neuroscience. Trends Neurosci. 38, 571–578 (2015).

44. D. Ongür, J. L. Price, The organization of networks within the orbital and medial prefrontal cortex of rats, monkeys and humans. Cereb. Cortex 10, 206–219 (2000).

45. Z. Samara, et al., Human orbital and anterior medial prefrontal cortex: Intrinsic connectivity parcellation and functional organization. Brain Struct. Funct. 222, 2941–2960 (2017).

46. J. O’Doherty, M. L. Kringelbach, E. T. Rolls, J. Hornak, C. Andrews, Abstract reward and punishment representations in the human orbitofrontal cortex. Nat. Neurosci. 4, 95–102 (2001).

47. E. T. Rolls, The orbitofrontal cortex and emotion in health and disease, including depression. Neuropsychologia 128, 14–43 (2019).

48. F. Grabenhorst, E. T. Rolls, B. A. Parris, From affective value to decision-making in the prefrontal cortex. Eur. J. Neurosci. 28, 1930–1939 (2008).

49. E. K. Miller, J. D. Cohen, An integrative theory of prefrontal cortex function. Annu. Rev. Neurosci. 24, 167–202 (2001).

50. D. H. Zald, et al., Meta-analytic connectivity modeling reveals differential functional connectivity of the medial and lateral orbitofrontal cortex. Cereb. Cortex 24, 232–248 (2014).

51. C. S. Carter, et al., Anterior cingulate cortex, error detection, and the online monitoring of performance. Science 280, 747–749 (1998).

52. S. N. Haber, H. Liu, J. Seidlitz, E. Bullmore, Prefrontal connectomics: from anatomy to human imaging. Neuropsychopharmacology 47, 20–40 (2022).

53. L. Pessoa, R. Adolphs, Emotion processing and the amygdala: from a “low road” to “many roads” of evaluating biological significance. Nat. Rev. Neurosci. 11, 773–783 (2010).

54. K. J. Ressler, Amygdala activity, fear, and anxiety: modulation by stress. Biol. Psychiatry 67, 1117–1119 (2010).

55. A. P. Maurer, L. Nadel, The Continuity of Context: A Role for the Hippocampus. Trends Cogn. Sci. 25, 187–199 (2021).

56. D. G. Weissman, et al., Earlier adolescent substance use onset predicts stronger connectivity between reward and cognitive control brain networks. Dev. Cogn. Neurosci. 16, 121–129 (2015).

57. A. C. K. van Duijvenvoorde, M. Achterberg, B. R. Braams, S. Peters, E. A. Crone, Testing a dual-systems model of adolescent brain development using resting-state connectivity analyses. Neuroimage 124, 409– 420 (2016).

58. J. Jung, M. A. Lambon Ralph, R. L. Jackson, Subregions of DLPFC Display Graded yet Distinct Structural and Functional Connectivity. J. Neurosci. 42, 3241–3252 (2022).

59. S. Berboth and C. Morawetz, Amygdala-prefrontal connectivity during emotion regulation: A meta-analysis of psychophysiological interactions. Neuropsychologia 153, 107767 (2021).

60. J. L. Robinson, et al., The functional connectivity of the human caudate: an application of meta-analytic connectivity modeling with behavioral filtering. Neuroimage 60, 117–129 (2012).

61. K. Jarbo, T. D. Verstynen, Converging structural and functional connectivity of orbitofrontal, dorsolateral prefrontal, and posterior parietal cortex in the human striatum. J. Neurosci. 35, 3865–3878 (2015).

62. K. Yuan, et al., The left dorsolateral prefrontal cortex and caudate pathway: New evidence for cue-induced craving of smokers. Hum. Brain Mapp. 38, 4644–4656 (2017).

63. B. Rao, et al., Development of functional connectivity within and among the resting-state networks in anesthetized rhesus monkeys. Neuroimage 242, 118473 (2021).

64. M. C. Stevens, The contributions of resting state and task-based functional connectivity studies to our understanding of adolescent brain network maturation. Neurosci. Biobehav. Rev. 70, 13–32 (2016).

65. J. H. Decker, A. R. Otto, N. D. Daw, C. A. Hartley, From Creatures of Habit to Goal-Directed Learners: Tracking the Developmental Emergence of Model-Based Reinforcement Learning. Psychol. Sci. 27, 848– 858 (2016).

66. V. P. Murty, F. Calabro, B. Luna, The role of experience in adolescent cognitive development: Integration of executive, memory, and mesolimbic systems. Neurosci. Biobehav. Rev. 70, 46–58 (2016).

67. F. J. Calabro, V. P. Murty, M. Jalbrzikowski, B. Tervo-Clemmens, B. Luna, Development of Hippocampal–Prefrontal Cortex Interactions through Adolescence. Cereb. Cortex 30, 1548–1558 (2019).

68. X. Song, et al., Strengths and challenges of longitudinal non-human primate neuroimaging. Neuroimage 236, 118009 (2021).

