## Supplemental_material_Deshpande_Kohut for "Age-related development in prefrontal-subcortical resting-state functional connectivity in nonhuman primates"

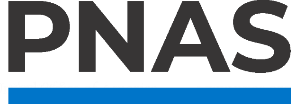

**Supporting Information for**

Age-related development in prefrontal-subcortical resting-state functional connectivity in nonhuman primates

Harshawardhan U. Deshpande and Stephen J. Kohut

Behavioral Neuroimaging Laboratory, McLean Hospital and Department of Psychiatry, Harvard Medical School, Belmont, MA, USA

Correspondence:

Harshawardhan U. Deshpande, Ph.D.

Stephen J. Kohut, Ph. D.

**This PDF file includes:**

Supporting text

Figures SF1 to SF2

Tables ST1 to ST2

SI References

**Supplementary methods:**

*Acclimation for Imaging Procedures.* Acclimation to the MRI apparatus typically occurred 5 days per week. Briefly, subjects were gradually trained to rest in a prone position on their haunches within a custom designed 3D printed (ABS plastic) chair enclosure with a 3D printed helmet mounted to the chair body with plastic screws. The helmet was lined with padding to limit motion and optimize comfort; sounds recorded from the scanner were played within the mock bore. The helmet included a platform to position a transmit/receive surface coil for data capture. Once subjects were acclimated to the mock MRI procedures, scan sessions began. Vital signs (i.e., heart rate, respiration rate, and oxygen saturation [SPO2]) were tracked and recorded at 5-min intervals throughout both training and MR sessions. For further details, please refer (1) and (2).

*MRI data acquisition.* Images were acquired with a 9.4 Tesla/400 mm diameter MR system (Varian Direct Drive, Varian Inc, Palo Alto, CA) within an 11.9 cm gradient bore (Resonance Research Inc, Billerica, MA). Preparatory scans included automated image-based shimming <75 Hz and fMRI images were acquired with a whole-brain gradient-echo planar imaging (EPI) sequence with TE=8 ms, TR=1500 ms, and flip angle=90°; scan matrix was 64 on a 64 mm field of view with 54 x 1 mm slices; scan time for 1200 volumes was 30 min. Distortion-matched anatomic images were acquired with a spin-echo EPI sequence (TE=17.5 ms, TR=1500 ms, flip angle =90°, averages=8; scan matrix=64 on a 64 mm field of view with 54 x 1 mm coronal slices matched to the fMRI slices).

*MRI data processing.* Data processing was performed using FSL and AFNI (23). The data were visually checked slice-by-slice for artifacts using all three orthogonal directions. Quality control was evaluated using MRI Quality Control tool (MRIQC; (24)), and group mean motion statistics were calculated for the data set. FMRIB’s Software Library (FSL, Oxford University, UK) image preprocessing pipeline was used to process the data as follows (25, 26): after slice-timing correction, the first 10 volumes from each scan were removed to allow for data stabilization. MCFLIRT tool in FSL (27) was used for head motion correction by volume realignment to the middle volume. Session averaged functional volumes were aligned to the VALiDATe (28) T1w template through a 12 DOF affine transformation followed by adjustment of nonlinear distortion fields using the jip analysis toolkit ([www.nitrc.org/projects/jip](http://www.nitrc.org/projects/jip)) for registration and brain extraction. Spatial smoothing was conducted using a Gaussian kernel of 2.0 mm FWHM. Temporal filtering was applied using a band pass filter (0.01 to 0.2 Hz, *fslmaths -bptf*).

*Minimization of head motion.* To minimize in-scanner head motion, subjects were extensively acclimated to awake scanning procedures as described above. A strict threshold of mean framewise displacement (FD)=0.3 mm across a session was used to exclude data; if a subject’s scan was excluded, they were rescanned within a week of the previous scan. During processing, intensity spiking was analyzed and adjusted for using an in-house program, “spikefix”, which is designed to find and remove spikes from fMRI datasets (<https://github.com/bbfrederick/spikefix>) with a threshold of 1 mm framewise displacement (30). Z-scored transforms of twelve motion parameters (three rotational, three translational, six first-order derivatives) were regressed from the data. Finally, average FD values for individual subject scans were used as nuisance covariates in subsequent analyses.

**Supplementary Results**

Seed-to-whole-brain FC differences between adolescents and adults for non-hub regions

| **#Voxels** | **Peak X** | **Peak Y** | **Peak Z** | **t-value** | **Peak region** |
| --- | --- | --- | --- | --- | --- |
| **R_vlPFC** | | | | | |
| 3783 | -2.3 | -3.3 | 11.8 | -5.75 | PCC |
| 190 | -0.3 | 17.6 | -7.9 | 4.72 | Cerebellum (medial) |
| 90 | -0.3 | -12.2 | 0.6 | 5.6 | Medial septum |
| 74 | 10.6 | 12.6 | -10.7 | 3.93 | Cerebellum (lateral) |
| 23 | -8.2 | 22.6 | -3.2 | 4.07 | R Occipital (V1) |
| **L_vlPFC** | | | | | |
| 4367 | -0.3 | -6.3 | 14.6 | -5.81 | Cingulate cortex |
| 457 | 0.7 | 4.7 | -10.7 | 4.51 | Nucleus Fasciculus Cuneatum |
| 239 | -0.3 | -12.2 | 0.6 | 6.21 | Medial septum |
| **lat OFC** | | | | | |
| 404 | -1.3 | 2.7 | 5.2 | -5.53 | PCC |
| 146 | -0.3 | -9.2 | -0.4 | 6.3 | Globus Pallidus |
| 31 | -9.2 | 4.7 | 2.4 | -4.12 | R Hippocampus |
| 28 | 5.7 | 3.7 | 9.9 | -4.42 | Intraparietal area |
| 23 | -0.3 | 4.7 | -11.6 | 3.64 | Nervus Ascuticus |
| 20 | -16.2 | -7.3 | -3.2 | -4.36 | Inf temporal gyrus |
| **vmPFC** | | | | | |
| 194 | 1.7 | 1.7 | 4.3 | -4.84 | Posterior cingulate |
| 27 | 6.7 | 2.7 | -3.2 | -3.81 | L lateral geniculate |
| 20 | -15.2 | -3.3 | -1.3 | -3.78 | R superior temporal |

**Table ST1. Seed-whole-brain functional connectivity differences for non-hub prefrontal ROIs.** For the ROIs, the brain regions as different between adolescents and adults identified using the squirrel monkey atlas are listed. The X, Y, Z coordinates are listed for the peak region.

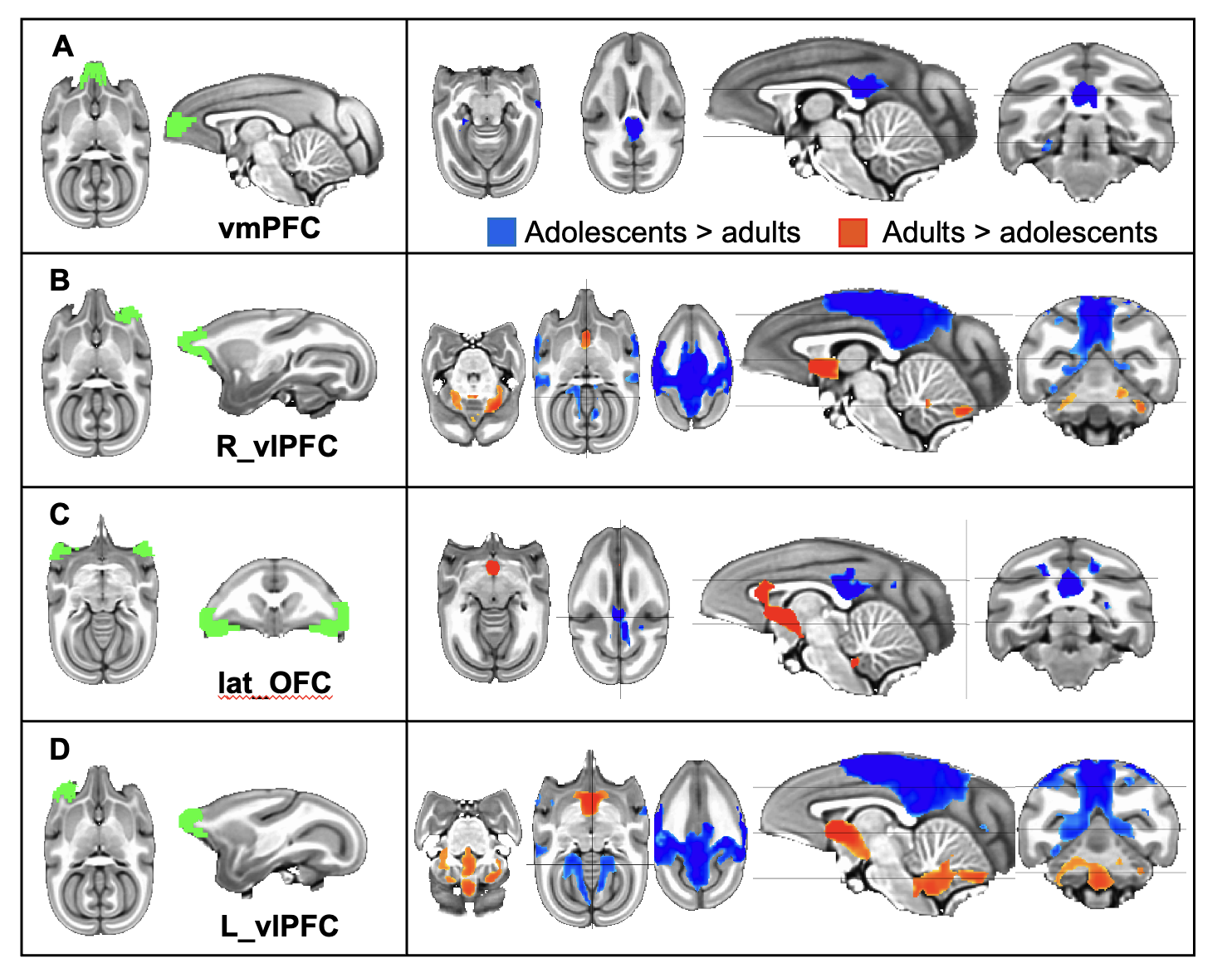

**Figure SF1. Seed-to-whole-brain functional connectivity differences between the adolescent and adult groups for non-hub regions.** p=0.001, FDR corrected p<0.05, minimum cluster threshold > 19 voxels

| **#Voxels** | **Peak X** | **Peak Y** | **Peak Z** | **t-value** | **Peak region** |
| --- | --- | --- | --- | --- | --- |
| **L Amygdala** | | | | | |
| 165 | 0.7 | -9.2 | -0.4 | 5.98 | Medial septum |
| 137 | 17.6 | -3.3 | -0.4 | 3.9 | L Temporalis superior cortex |
| 80 | -11.2 | 15.6 | 6.2 | 4.71 | R Occipital Cortex (V1) |
| 36 | -7.3 | -3.3 | -5 | 4.86 | R Hippocampus |
| 33 | -16.2 | -3.3 | 1.5 | 3.7 | R Temporalis superior cortex |
| **R Caudate** | | | | | |
| 348 | -2.3 | -4.3 | 8 | -3.92 | R Cingulate |
| 189 | -10.2 | -3.3 | 7.1 | -4.23 | R Ant Insula |
| 55 | 3.7 | 3.7 | -2.2 | -4.29 | Commisura coliculi inferioris |
| 46 | -7.3 | -0.3 | -5 | -4.09 | R Caudate nucleus |
| 34 | 5.7 | -7.3 | -0.4 | -4.78 | Thalamus |
| 31 | -7.3 | -6.3 | 3.4 | -3.61 | R Putamen |
| **L vStr** | | | | | |
| 832 | 8.6 | 14.6 | 2.4 | 6.98 | L visual cortex |
| 667 | -7.3 | 18.6 | 4.3 | 5.11 | R visual cortex |
| 105 | 9.6 | -21.2 | 5.2 | 7.66 | L vlPFC |
| 61 | 2.7 | -6.3 | -6.9 | -4.88 | Chiasma nervorum opticorum |
| 45 | -9.2 | -11.2 | -10.7 | 3.89 | R Amygdala |
| 43 | 1.7 | -6.3 | 6.2 | -4.65 | Cingulate |
| 31 | 0.7 | 1.7 | 3.4 | -4.31 | Fornix |
| 27 | 4.7 | -2.3 | -8.8 | -4.25 | Pedunculis medius cerebelli |
| 22 | -14.2 | -13.2 | 0.6 | 3.69 | R Inf Frontal gyrus |
| **L Putamen (p=0.001, FDR q=0.055)** | | | | | |
| 47 | -3.3 | -4.3 | 9 | -4.47 | R Cingulate |
| 44 | -9.2 | -3.3 | 6.2 | -5.41 | R Ant Insula |
| 39 | 9.6 | -0.3 | 5.2 | -4.18 | L Pos Insula |
| 33 | -2.3 | 3.7 | 3.4 | -5.29 | Cingulate gyrus |
| 21 | -12.2 | 14.6 | 0.6 | 3.9 | R Occipital cortex (V1) |
| **R vStr** | | | | | |
| 595 | 9.6 | 14.6 | 3.4 | 5.85 | L visual cortex |
| 513 | -8.2 | 17.6 | 3.4 | 5.26 | R visual cortex |
| 66 | -1.3 | -7.3 | -7.9 | -4.64 | Pes pedunculi |
| 61 | 8.6 | -21.2 | 6.2 | 6.03 | L vlPFC |
| 29 | -4.3 | 16.6 | 8 | 3.93 | R visual cortex |
| 28 | 0.7 | 2.7 | 3.4 | -5.08 | Fornix |
| **Continued on the next page** | | | | | |
| **#Voxels** | **Peakx** | **Peaky** | **Peakz** | **t-value** | **Peak region** |
| **R Putamen (p=0.001, FDR q=0.12)** | | | | | |
| 61 | -3.3 | -4.3 | 9.0 | -4.97 | R Middle Cingulate |
| 60 | -1.3 | 1.7 | 3.4 | -4.39 | Corpus Callosum |
| **R Hippocampus (p=0.001, FDR q=0.058)** | | | | | |
| 124 | -8.2 | 15.6 | 4.3 | 4.27 | R visual cortex |
| 65 | 9.6 | 15.6 | 3.4 | 3.69 | L visual cortex |
| 63 | -1.3 | -7.3 | -7.9 | -5.27 | Medial hypothalamus |
| 45 | 0.7 | -7.3 | 5.2 | -4.94 | Caudate |

**Table ST2. Seed-whole-brain functional connectivity differences for non-hub subcortical ROIs.** For the ROIs, the brain regions as different between adolescents and adults identified using the squirrel monkey atlas are listed. The X, Y, Z coordinates are listed for the peak region.

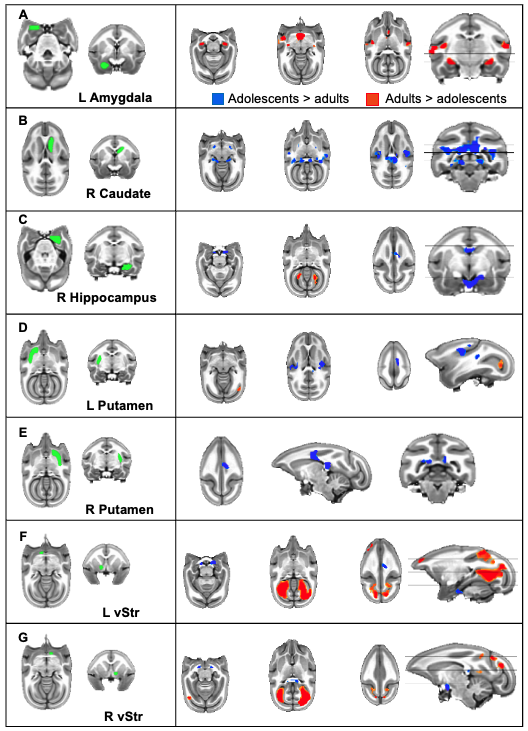

**Figure SF2. Seed-to-whole-brain functional connectivity differences between the adolescent and adult groups for non-hub regions.** p=0.001, minimum cluster threshold > 19 voxels

**REFERENCES**

1. W. Yassin, *et al.*, Resting state networks of awake adolescent and adult squirrel monkeys using ultra-high field (9.4T) functional magnetic resonance imaging. *bioRxiv* (2023) https:/doi.org/[10.1101/2023.01.08.523000](http://dx.doi.org/10.1101/2023.01.08.523000).

2. S. L. Withey, *et al.*, Fentanyl-induced changes in brain activity in awake nonhuman primates at 9.4 Tesla. *Brain Imaging Behav.* **16**, 1684–1694 (2022).

3. L. Cao, S. J. Kohut, B. D. Frederick, Estimating and mitigating the effects of systemic low frequency oscillations (sLFO) on resting state networks in awake non-human primates using time lag dependent methodology. *Front. Neuroimaging* **1**, 1031991 (2023).

4. O. Esteban, *et al.*, MRIQC: Advancing the automatic prediction of image quality in MRI from unseen sites. *PLoS One* **12**, e0184661 (2017).

5. S. M. Smith, *et al.*, Advances in functional and structural MR image analysis and implementation as FSL. *Neuroimage* **23 Suppl 1**, S208–19 (2004).

6. M. Jenkinson, C. F. Beckmann, T. E. J. Behrens, M. W. Woolrich, S. M. Smith, FSL. *Neuroimage* **62**, 782–790 (2012).

7. M. Jenkinson, P. Bannister, M. Brady, S. Smith, Improved optimization for the robust and accurate linear registration and motion correction of brain images. *Neuroimage* **17**, 825–841 (2002).

8. K. G. Schilling, *et al.*, The VALiDATe29 MRI Based Multi-Channel Atlas of the Squirrel Monkey Brain. *Neuroinformatics* **15**, 321–331 (2017).
